# Activin A targets extrasynaptic NMDA receptors to improve neuronal and behavioral deficits in a mouse model of Huntington disease

**DOI:** 10.1101/2023.09.06.556580

**Authors:** Wissam B. Nassrallah, Daniel Ramandi, Judy Cheng, Jean Oh, James Mackay, Marja D. Sepers, David Lau, Hilmar Bading, Lynn A. Raymond

## Abstract

Cortical-striatal synaptic dysfunction, including enhanced toxic signaling by extrasynaptic N-methyl-D-aspartate receptors (eNMDARs), precedes neurodegeneration in Huntington disease (HD). A previous study showed Activin A, whose transcription is upregulated by calcium influx via synaptic NMDARs, suppresses eNMDAR signaling. Therefore, we examined the role of Activin A in the YAC128 HD mouse model, comparing it to wild-type controls. We found decreased Activin A secretion in YAC128 cortical-striatal co-cultures, while Activin A overexpression in this model rescued altered eNMDAR expression. Striatal overexpression of Activin A *in vivo* improved motor learning on the rotarod task, and normalized striatal neuronal eNMDAR-mediated currents, membrane capacitance and spontaneous excitatory postsynaptic current frequency in the YAC128 mice. These results support the therapeutic potential of Activin A signaling and targeting eNMDARs to restore striatal neuronal health and ameliorate behavioral deficits in HD.

## Introduction

Huntington Disease (HD) is a fatal hereditary neurodegenerative disorder characterized by a movement disorder including chorea or involuntary dance-like movements, dystonia, bradykinesia, and gait disturbances(Ghosh and Tabrizi, 2018). HD patients also often experience cognitive changes such as decreased processing speed and impaired executive and visuospatial function before the onset of motor symptoms (Paulsen et al., 2014; Tabrizi et al., 2013). HD is caused by a CAG trinucleotide repeat expansion in the gene coding the protein huntingtin (HTT), resulting in an expanded polyglutamine tract near the N-terminus of this protein. More than 35 repeats lead to HD, and the age of onset is inversely proportional to the number of CAG repeats(MacDonald et al., 1993). At the cellular level, HD is characterized predominantly by degeneration of striatal medium-sized spiny neurons (MSNs), as well as the loss of cortical pyramidal neurons and certain classes of interneurons to a lesser degree (Plotkin and Surmeier, 2015; Raymond et al., 2011).

One reported mechanism that can contribute to striatal neuronal dysfunction and death in HD involves aberrant signaling by glutamate-gated ionotropic N-methyl-D-aspartate receptors (NMDAR) via a relative increase in activation of extrasynaptic versus synaptic NMDAR. Calcium influx via synaptic NMDAR stimulates neuroprotective signaling pathways required for synaptogenesis, synaptic plasticity involved in learning and memory, and activation of pro-survival genes that maintain neuronal health through induction of cyclic-AMP response element-binding protein (CREB) phosphorylation and brain-derived neurotrophic factor (BDNF) expression. In contrast, extrasynaptic NMDAR (eNMDAR) activation due to excessive glutamate release and spillover outside synapses opposes plasticity and promotes cell death pathways in a dominant manner(Hardingham and Bading, 2010). Increased eNMDAR signaling and decreased BDNF expression, leading to synaptic dysfunction and learning impairments, has been documented in other neurodegenerative diseases as well(Parsons and Raymond, 2014). Impaired NMDAR-dependent plasticity and augmented eNMDAR signaling has been reported in HD transgenic mice prior to manifestation of motor symptoms(Milnerwood et al., 2010a; Plotkin et al., 2014). Therefore, there is a strong rationale to pharmacologically inhibit eNMDARs in HD and other neurodegenerative diseases. However, selectively blocking eNMDARs without off-target effects has been difficult, and given the wide involvement of NMDARs in many processes, drugs that inhibit both synaptic and extrasynaptic NMDARs can produce serious side effects(Lipton, 2006; Xia et al., 2010).

Activin A, the dimerized form of Inhibin beta A (INHBA), has been shown to be an important neurotrophic factor that regulates MSN differentiation, dendritic spine morphology, and glutamatergic neurotransmission(Su et al., 2018). Furthermore, Activin A is transcribed and secreted following synaptic activity in a nuclear-calcium-dependent manner, and reduces toxic eNMDAR signaling through unknown mechanisms(Lau et al., 2015). We hypothesized that the synaptic dysfunction and reduced BDNF levels observed in HD lead to decreased INHBA signaling, contributing to increased eNMDAR activation.

Our findings reveal that cortical-striatal co-cultures from HD transgenic mice exhibit reduced Activin A secretion, and that INHBA overexpression in these cultures normalized the balance of synaptic and extrasynaptic NMDAR expression. Notably, we validated these results *in vivo,* showing that Activin A overexpression in the striatum of HD transgenic mice alleviated motor learning impairments, reduced striatal neuronal eNMDAR currents, and normalized other measures of striatal neuronal health.

## Results

### Activin A secretion is decreased in YAC128 cortical-striatal co-cultures

Excessive eNMDAR activation has been shown previously to reduce transcription of a group of genes involved in promoting neuronal health and survival, including Activin A(Hardingham and Bading, 2010). We began by measuring Activin A levels in WT and YAC128 mouse-derived cortical-striatal co-cultures, a preparation in which YAC128 striatal neurons show a robust increase in cell-surface GluN2B-containing eNMDAR expression(Botelho et al., 2014; Kovalenko et al., 2018; Milnerwood et al., 2010b; Plotkin et al., 2014). Given that Activin A is released in the extracellular space, it can be measured from the culture media. To minimise variability in week-to-week culture handling, WT and YAC128 cultures were plated the same day and Activin A levels measured across different(Hardingham and Bading, 2010) days *in vitro* (DIV) were normalised to levels measured at DIV4 within each culture batch. Although both genotypes showed a robust increase in Activin A secretion at DIV21 relative to DIV4, this increase was significantly smaller in YAC128 culture media compared to that of WT cultures (Fig. 1A). To explore whether this difference is mainly derived from the cortical or striatal neurons, we used chimeric co-cultures, combining either WT or YAC128 cortical cells with either WT or YAC128 striatal cells (4 combinations). Our findings showed that the decrease in Activin A secretion was only seen when the cortical neurons were derived from YAC128 mice, regardless of the genotype of the striatal neurons (Fig. 1B). This suggests that the cortical neurons mediate diminished Activin A secretion observed in YAC128 co-cultures.

**Figure 1.**
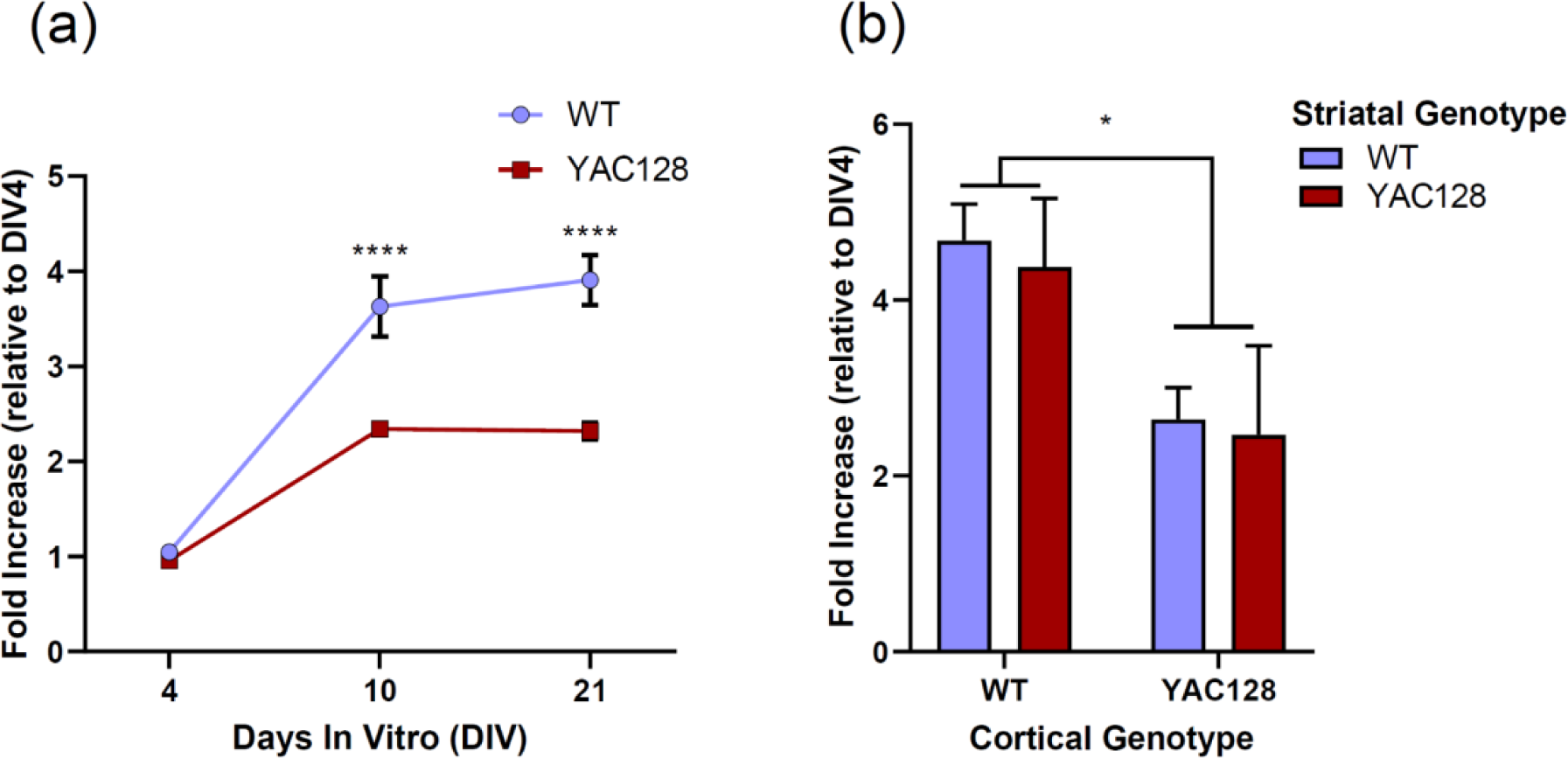
Activin A secretion is decreased in YAC128 cortical-striatal co-cultures. ***a)*** Activin A concentration average in WT and YAC128 culture media presented as a fold increase at DIV21 and DIV10 and DIV4 compared to DIV4. Data were analysed with a two-way ANOVA (p-values: genotype = <0.0001, DIV = <0.0001, interaction = 0.0005). (WT, n = 5 cultures, YAC128, n = 5 cultures). ***b*)** Activin A concentration average in chimeric co-culture media (combining either WT or YAC128 cortical cells with either WT or YAC128 striatal cells) presented as a fold increase at DIV21 compared to DIV4. Data were analysed with a two-way ANOVA (p-values: cortical genotype = 0.0151, striatal genotype = 0.7365, interaction = 0.9291; n=4 measurements for each DIV, from 2 independent culture batches, for each of the four chimeric culture combinations). DIV = days *in vitro*. Data are represented as mean ± SEM (WT = Wild-type).

Previous research has shown that the expression and release of Activin A requires pre-synaptic glutamate and BDNF release, as well as activation of postsynaptic NMDA receptors(Lau et al., 2015). Reduced cortical neuronal activity and BDNF release has been reported in primary neuronal culture and tissue from HD mice(Lenoir et al., 2022; Mackay et al., 2023; Virlogeux et al., 2018; Zuccato and Cattaneo, 2009), and this could lead to a decrease in Activin A expression and release in MSNs, highlighting the importance of cortical neurons in Activin A secretion. Two strategies to correct this impairment in the MSNs could be to overexpress Activin A in the striatum or to correct putative changes in cortical activity and/or BDNF release. In this study, we pursued the former approach.

### Activin A overexpression normalizes GluN2B surface expression in YAC128 striatal neurons

Activin A overexpression in hippocampal neurons has been shown to reduce calcium influx mediated by eNMDARs, although the mechanism remains unclear(Lau et al., 2015). In HD mice, surface expression of GluN2B-containing NMDARs is increased at extrasynaptic sites in striatal neurons, leading to increased vulnerability to MSN loss(Milnerwood et al., 2012, 2010b; Okamoto et al., 2009). Previously, we showed that the increased surface/internal ratio of GluN2B in HD cortical-striatal co-cultures is a reliable proxy for enhanced eNMDAR expression(Milnerwood et al., 2012), consistent with reports that GluN2B-containing NMDARs predominantly localize to extrasynaptic sites, whereas these receptors are internalized and/or expressed at lower levels in the synapse(Sanz-Clemente et al., 2013). Given Activin A’s ability to reduce toxic eNMDAR-mediated calcium influx(Lau et al., 2015), we assessed its effect on surface GluN2B expression. To do this, co-cultures overexpressing Activin A (using an rAAV strategy) were immuno-stained to image surface vs. internal GluN2B, as described previously(Milnerwood et al., 2012). Results show that Activin A overexpression in cortical-striatal co-cultures decreased the surface/internal GluN2B ratio in YAC128 MSNs back to that of WT MSNs, without changing the ratio in WT MSNs (Fig. 2), suggesting that increased Activin A levels normalize GluN2B-eNMDAR expression in striatal neurons.

**Figure 2.**
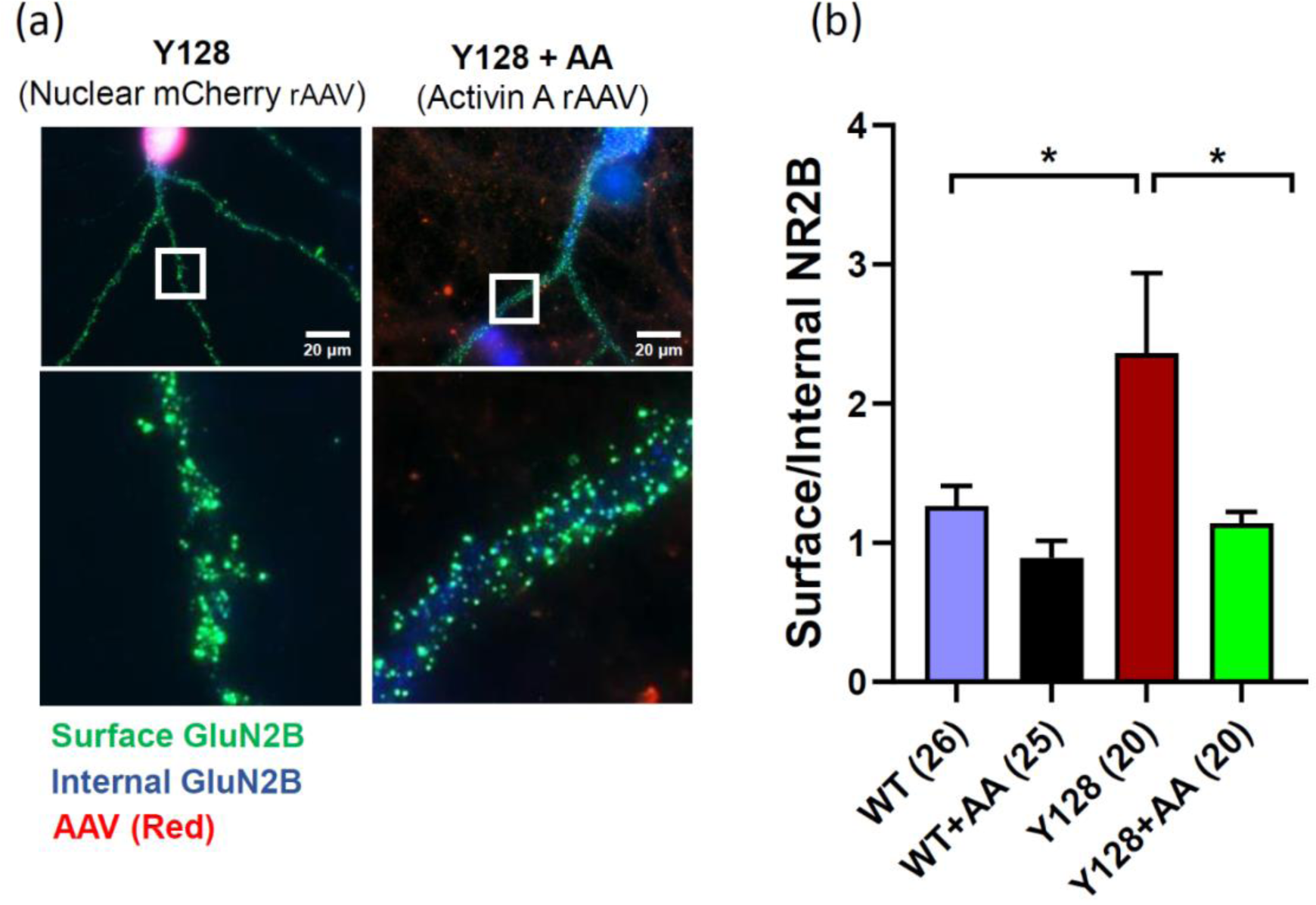
Activin A overexpression normalizes GluN2B surface expression in YAC128 striatal neurons. ***a)*** Example of MSN cells stained for internal (blue) and surface (green) GluN2B. Nucleus-localised mCherry rAAV (Left) and Activin A rAAV (Right) treated cells are in red. The rAAV was applied to the culture wells at DIV 7, and then stained at DIV21 as in the STAR methods section. ***b)*** Surface/internal GluN2B average of WT and YAC128 MSNs treated with either nucleus-localised mCherry rAAV or Activin A rAAV (AA). Data were analysed with a two-way ANOVA (p-values: genotype = 0.0099, treatment = 0.0009, interaction = 0.2950; Bonferroni multiple comparisons test: Control vs. Activin A: WT p-value = 0.1476; YAC128 p-value = 0.0075). (WT, n = 26 cells (4 cultures); WT Activin A, n = 25 cells (4 cultures); YAC128, n = 19 cells (4 cultures); YAC128 Activin A, n = 20 cells (4 cultures)). Data are represented as mean ± SEM (AA = Activin A, WT = Wild-type, Y128 = YAC128).

### Activin A overexpression increases the ability of YAC128 mice to learn the rotarod task

Previously, we showed that a 2-month treatment with low dose memantine, which preferentially blocks eNMDARs, improved motor learning on the rotarod task in 4-to 5-month-old YAC128 mice (Milnerwood et al., 2010b). To test whether Activin A overexpression can improve motor learning in YAC128 mice, 6-week-old male and female mice were injected with either an Activin A rAAV or a control mCherry rAAV in the striatum bilaterally. Pilot experiments confirmed robust Activin A/mCherry expression in striatum 2-3 weeks following injection of 0.5 or 1uL (Fig. 3A); since the latter resulted in more extensive coverage of the striatum, this higher dose was used for all experiments. Mice were then tested at 6 months of age using the rotarod motor learning task. The rotarod task measures motor coordination and balance, in which mice are placed on a rotating rod that accelerates in speed over 5 minutes, and the latency to fall is recorded over 4 consecutive days of 3 trials each to determine rate of learning and overall motor coordination. We found that YAC128 mice injected with the control rAAV performed significantly worse than WT mice across all 4 days, failing to show robust learning (Fig. 3B, C, D). In contrast, Activin A overexpression in YAC128 mice rescued the motor learning and coordination impairments to levels comparable to WT mice. Similar results were obtained in both male and female mice (Fig. 3 Supplement). Notably, this result was not dependent on the mouse’s weight (Fig. 3E), as Activin A overexpression did not have an effect on the well-known increased weight of YAC128 mice compared to WT mice (Pouladi et al., 2010; Van Raamsdonk et al., 2006). The rotarod performance of WT mice was not significantly affected by striatal overexpression of Activin A.

**Figure 3.**
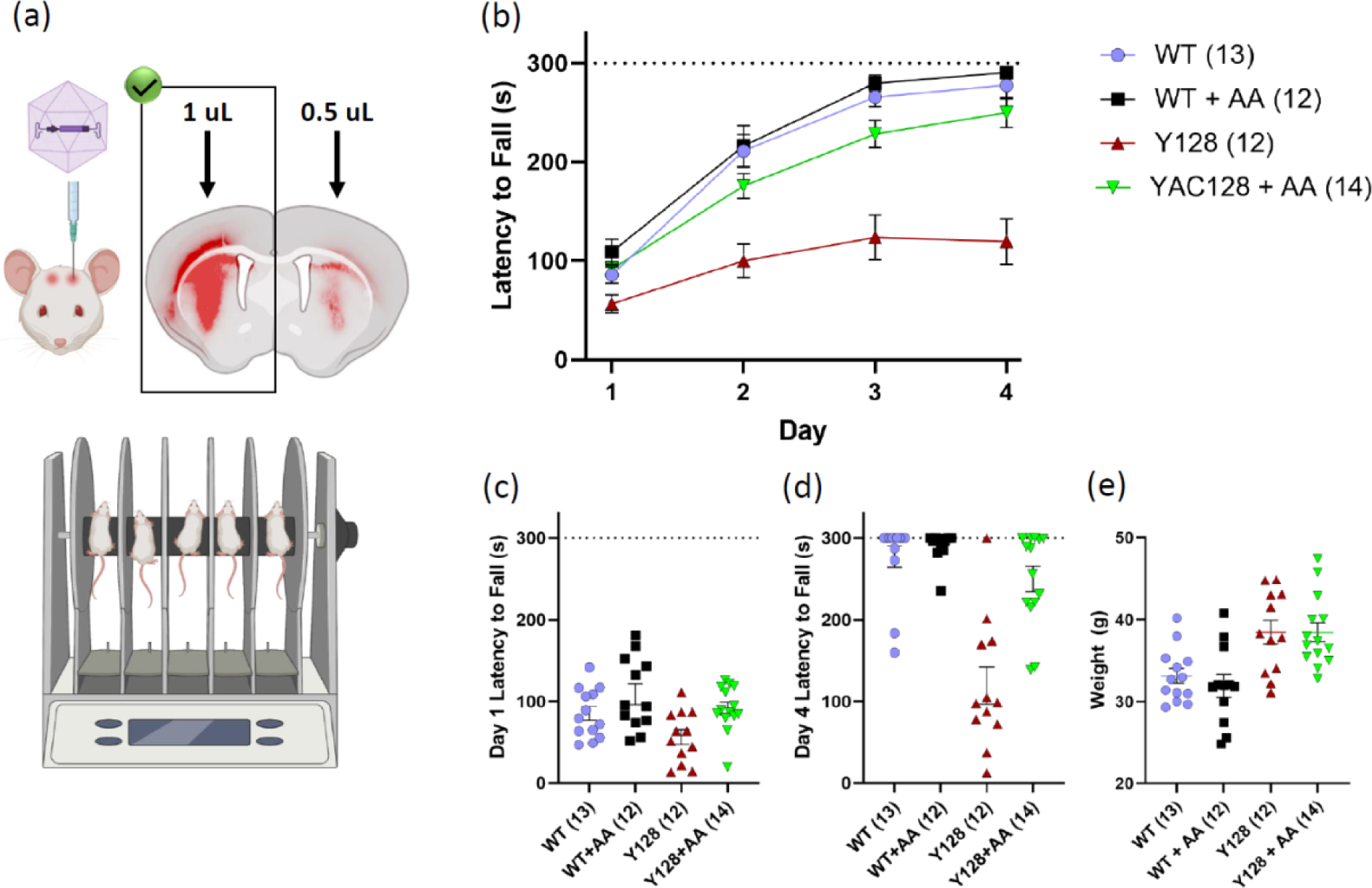
Activin A overexpression increases the ability of YAC128 mice to learn the rotarod task. ***a)*** Example of a coronal brain slice expressing Activin A (right hemisphere injected with 1 μL rAAV and left hemisphere injected with 0.5 μL rAAV) with a cartoon showing the injection procedure done at 1.5 months of age, and another cartoon showing the rotarod experiments done at 6 months of age. ***b)*** Latency to fall averages per day of WT and YAC128 male and female mice at 6 months of age treated either with Activin A rAAV or a control mCherry rAAV. Data were analysed with a two-way ANOVA (p-values: genotype = <0.0001, Days = <0.0001, interaction = <0.0001). Number in parentheses indicates number of animals. ***c)*** Day 1 average latency to fall of WT and YAC128 male and female mice at 6 months of age treated either with Activin A rAAV or a control mCherry rAAV. Points represent individual mice. Data were analysed with Šídák’s multiple comparisons test (p-values: Y128 vs. WT = 0.8708, Y128 vs. WT+AA = 0.1420, Y128 vs. Y128+AA = 0.6384). ***d)*** Day 4 average latency to fall of WT and YAC128 male and female mice at 6 months of age treated either with Activin A rAAV or a control mCherry rAAV. Points represent individual mice. Data were analysed with Šídák’s multiple comparisons test (p-values: Y128 vs. WT = <0.0001, Y128 vs. WT+AA = <0.0001, Y128 vs. Y128+AA = <0.0001). ***e)*** Average weights of WT and YAC128 male and female mice at 6 months of age treated either with Activin A rAAV or a control mCherry rAAV. Points represent individual mice. Data were analysed with a two-way ANOVA (p-values: genotype = <0.0001, Treatment = 0.6161, interaction =0.6351). (WT, n = 13 mice; WT Activin A, n = 12 mice; YAC128, n = 12 mice; YAC128 Activin A, n = 14 mice). Data are represented as mean ± SEM (AA = Activin A, WT = Wild-type, Y128 = YAC128).

### Activin A overexpression normalizes the contribution of extrasynaptic NMDARs in YAC128 MSNs *ex vivo*

To investigate the effect of striatal Activin A overexpression on eNMDARs, voltage-clamped whole-cell patch recordings were made from MSNs in acute cortico-striatal brain slices within a month of the mice completing the rotarod task. DL-TBOA – a glutamate transporter inhibitor that promotes activation of eNMDARs through glutamate spillover – was used to assess differences in evoked NMDAR-mediated currents in MSNs. The area under the curve of current decay following the last stimulation in a 10-stimulation train (at 20 Hz) was normalized to the peak current amplitude of the 10^th^ response to determine the contribution of eNMDARs (Fig. 4A, B), which are enriched in GluN2B-containing NMDARs. The results showed that MSNs from control mCherry-injected YAC128 mice exhibited a larger TBOA-induced increase in area/peak compared to WTs (Fig. 4C, D), indicating a greater contribution from eNMDARs. However, in YAC128 mice with Activin A overexpression, this increase was normalized to WT levels, consistent with our findings in cortical-striatal co-cultures, where Activin A overexpression normalized the surface-to-internal ratio of GluN2B-containing NMDARs. These results suggest that Activin A overexpression can effectively down-regulate the eNMDAR-mediated currents in YAC128 MSNs.

**Figure 4.**
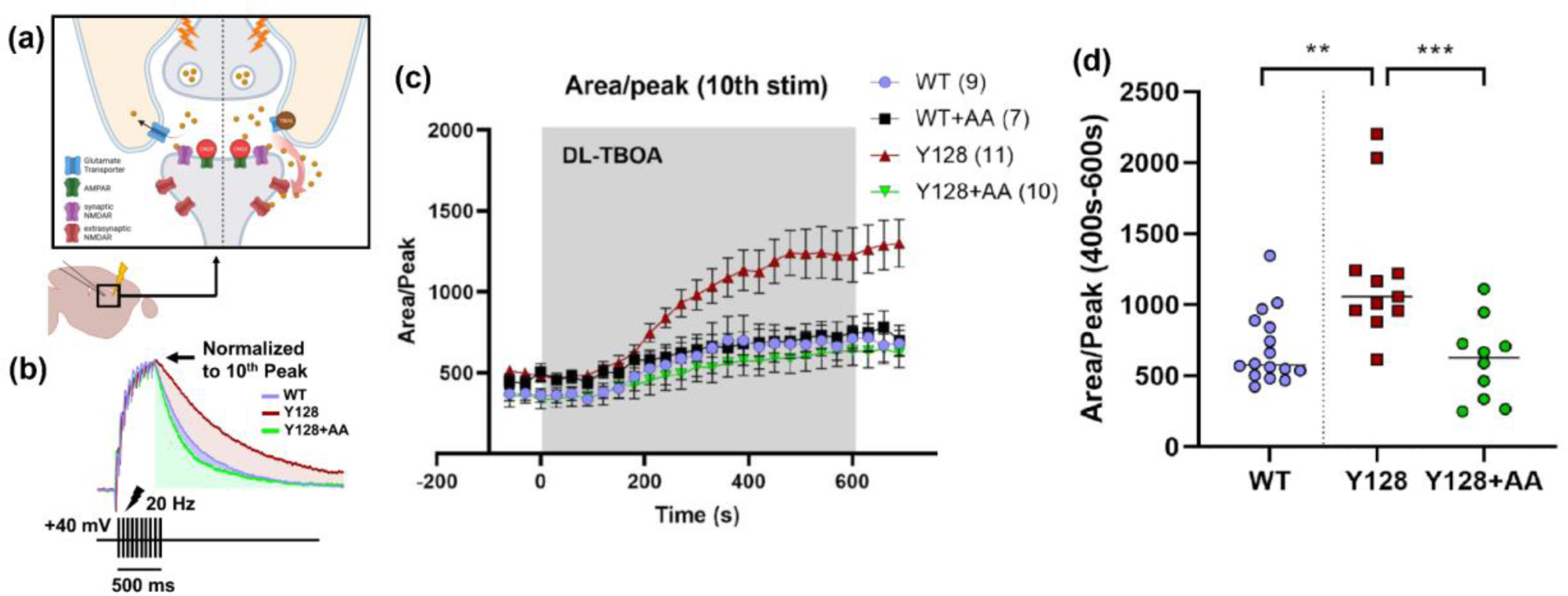
Activin A overexpression normalizes the contribution of extrasynaptic NMDARs in YAC128 MSNs *ex vivo*. ***a)*** A schematic depicting a synapse and the experimental paradigm to electrophysiologically study eNMDAR currents. To isolate NMDAR current, CNQX (10 μM, to block the AMPA receptors (green)), glycine (10 μM, NMDAR co-agonist), and LY341495 (1uM, to inhibit presynaptic metabotropic glutamate receptors) were utilised. DL-TBOA (10 μM), a glutamate transport inhibitor, was employed to cause glutamate spillover beyond the synapse into extrasynaptic sites, activating eNMDARs. Both synaptic (purple) and extrasynaptic (dark red) NMDARs are activated. The schematic also depicts that the recording electrode is in the striatum voltage clamping an MSN and a nearby stimulating electrode (thunderbolt) is also in the striatum of a sagittal brain slice. ***b)*** Example of voltage-clamp recording of three different MSNs showing their responses to stimulation train (20 Hz for 500 ms) during DL-TBOA in a WT MSN treated with a control mCherry rAAV (blue), YAC128 MSN treated with a control mCherry rAAV (red), and a YAC128 MSN treated with an Activin A rAAV (green). ***c)*** Time course of area/peak averages (every 30 seconds) of WT and YAC128 MSNs both previously treated with either Activin A rAAV or mCherry rAAV. Data were analysed with a two-way ANOVA (p-values: genotype = 0.0021, time = <0.0001, interaction = <0.0001). ***d)*** Area/peak averages from 400 to 600 seconds of WT and YAC128 MSNs previously treated with either Activin A rAAV or mCherry rAAV; data from the WT mCherry and Activin A treatments were pooled for averaging. Data were analysed using a two-way ANOVA (p-values: genotype = 0.0778, treatment = 0.0187, interaction = 0.0099; Šídák’s multiple comparisons test p-values: WT vs. Y128 = 0.0111, Y128 vs. Y128+AA = 0.0021, WT vs. Y128+AA = 0.9978). (WT, n = 9 cells; WT Activin A, n = 7 cells; YAC128, n = 11 cells; YAC128 Activin A, n = 10 cells). Data are represented as mean ± SEM (AA = Activin A, WT = Wild-type, Y128 = YAC128).

### MSN function and cell health is normalized in YAC128 mice overexpressing Activin A

We used electrophysiology to assess changes in the intrinsic membrane properties, basal excitatory synaptic transmission of striatal MSNs *ex vivo*, following termination of the rotarod experiments. Patch clamp recordings from acute brain slice showed that membrane capacitance, which increases in proportion to cell size in neurons, was significantly decreased in control YAC128 MSNs compared to WTs; however, in MSNs from YAC128 mice with Activin A overexpression, the capacitance was normalized to WT levels (Fig. 5A). We also recorded spontaneous excitatory postsynaptic currents (sEPSCs; Fig. 5C) and examined the frequency and amplitude of these currents as a measure of synaptic transmission. sEPSC frequency was found to be reduced in control YAC128 MSNs compared to WT MSNs, and was normalized in YAC128 mice with Activin A overexpression (Fig. 5D). Membrane resistance and sEPSC amplitude were not different between genotypes, nor were they impacted by Activin A expression (Fig. 5B, E). Taken together, these results suggest that Activin A overexpression has a beneficial effect on both neuronal function and health in the striatum of YAC128 mice.

**Figure 5.**
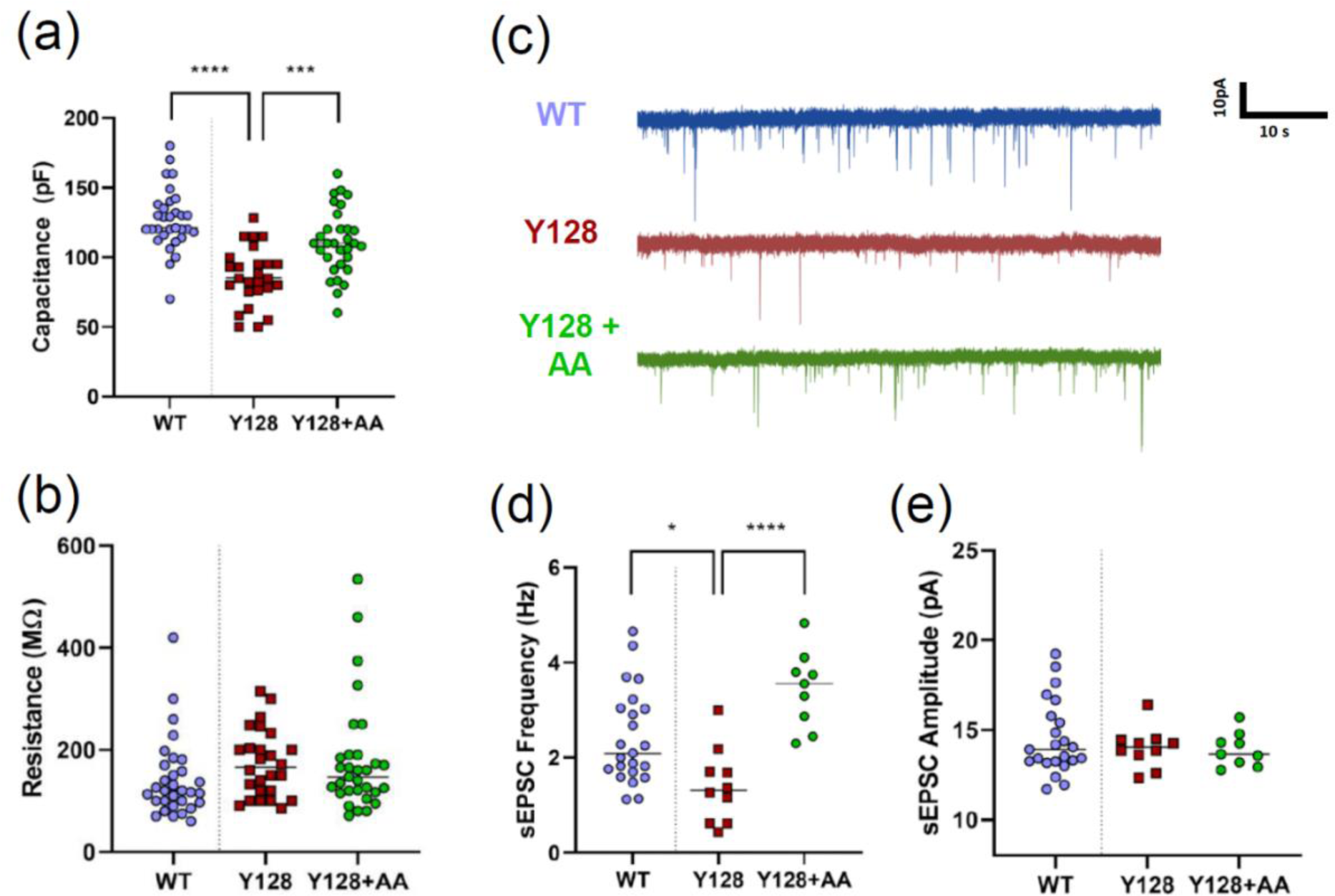
MSN function and cell health is normalized in YAC128 mice overexpressing Activin A. ***a)*** Cell capacitance averages of WT and YAC128 MSNs previously treated with either Activin A rAAV or mCherry rAAV. WT data from mCherry and Activin A treated mice were pooled. Data were analysed with a one-way ANOVA (p-value = <0.0001; Šídák’s multiple comparisons test p-values: WT vs. Y128 = <0.0001, Y128 vs. Y128+AA = 0.0002, WT vs. Y128+AA = 0.0149). ***b)*** Cell membrane resistance averages of WT and YAC128 MSNs previously treated with either Activin A rAAV or mCherry rAAV. Data were analysed with a one-way ANOVA (p-value = 0.2103). ***c)*** Example of electrophysiological traces showing sEPSCs in a WT + control mCherry MSN, YAC128 + control mCherry MSN, and YAC128 + Activin A MSN. ***d)*** sEPSCs frequency averages of WT and YAC128 MSNs previously treated with either Activin A rAAV or mCherry rAAV. Data were analysed with a one-way ANOVA (p-value = <0.0001; Šídák’s multiple comparisons test p-values: WT vs. Y128 = 0.0140, Y128 vs. Y128+AA = <0.0001, WT vs. Y128+AA = 0.0193). ***e)*** sEPSCs amplitude averages of WT and YAC128 MSNs previously treated with either Activin A rAAV or mCherry rAAV. Data were analysed with a one-way ANOVA (p-value = 0.5697). WT data from mCherry and Activin A treated mice were pooled for analysis in ***a – e***. (***a-b***: WT, n = 31 cells; YAC128, n = 26 cells; YAC128 Activin A, n = 31 cells). (***d-e***: WT, n = 25 cells; YAC128, n = 11 cells; YAC128 Activin A, n = 9 cells). Data are represented as mean ± SEM (AA = Activin A, WT = Wild-type, Y128 = YAC128).

## Discussion

In the present study, we have provided novel insights into the involvement of Activin A in the pathophysiology of HD, with a particular emphasis on its effect on striatal eNMDAR activity and expression, striatal neuronal health, and motor learning in the YAC128 HD mouse model. Compared with WT controls, our results demonstrated a significant reduction in Activin A secretion in YAC128 cortical-striatal co-cultures, with the primary contribution stemming from the cortical neurons. Importantly, we found that overexpressing Activin A in YAC128 effectively normalized the surface-to-internal GluN2B ratio – indicative of eNMDAR expression – of striatal neurons in cortical-striatal co-culture, and led to a marked decrease in striatal neuronal eNMDAR currents in brain slice, restoring them to levels observed in WT mice. Furthermore, our data revealed that early striatal overexpression of Activin A in YAC128 mice reversed the well-documented decline in motor learning performance and normalized markers of reduced striatal neuronal health typically associated with HD progression. Together, these findings underscore the potential significance of our study in the development of innovative therapeutic approaches targeting Activin A and its regulation of eNMDAR signaling, which may ultimately contribute to alleviating symptoms and slowing progression of HD.

### Activin A secretion is reduced and eNMDAR expression is increased in YAC128 co-cultures

Our *in vitro* experiments focused on the reduced secretion of Activin A and increased striatal neuron eNMDAR expression in YAC128 co-cultures. We observed that Activin A secretion is reduced in co-cultures containing YAC128 cortical neurons compared to WT cultures. Activin A, a key component in differentiating human pluripotent stem cells into MSNs(Arber et al., 2015), is essential for the healthy maturation and maintenance of cell-type identity in striatal neurons. Upon overexpressing Activin A in both cortical and striatal neurons, we found that the surface-to-internal GluN2B ratio in MSNs was normalized. Together, our results suggest that normal cortical activity and/or BDNF release promote Activin A secretion, which directly interacts with MSNs. These findings are consistent with previous HD research that suggests cortical activity drives MSN activity and health(Estrada-Sánchez et al., 2015; Virlogeux et al., 2018).

A normal level of Activin A release is not the only factor regulating eNMDARs, however. While we did not measure eNMDARs in the chimeric cultures in this study, previous research has shown that in co-cultures of WT cortex and YAC128 striatum, the striatal eNMDAR proportion remains higher than in pure WT co-cultures(Milnerwood et al., 2012). This indicates that both healthy cortex and healthy MSNs are necessary for the correct proportion of striatal eNMDARs. Taken together, the absence of mutant Huntingtin protein (mHTT) in cortical neurons is sufficient to normalize Activin A levels in cortical-striatal co-cultures, but only overexpression of Activin A can rescue the balance of synaptic to eNMDAR in YAC128 MSNs.

### Overexpression of Activin A improves motor learning and performance on the rotarod task in YAC128 mice

In our *in vivo* study, we observed remarkable improvements in motor learning and performance among YAC128 mice with Activin A overexpression. The accelerating rotarod task serves as a reliable measure of motor learning, and our findings showed that both male and female YAC128 mice with Activin A overexpression exhibited performance levels on par with their WT counterparts. Importantly, WT mice were not adversely affected by Activin A overexpression, suggesting that Activin A possesses low toxicity and might be a safe treatment option.

The improvements in motor learning and performance observed in YAC128 mice overexpressing Activin A could be attributed, in part, to two pathways. The first pathway involves Activin A’s role as a growth factor in promoting cellular health and preventing MSN dedifferentiation. Activin A facilitates differentiation of MSNs from human pluripotent stem cells(Arber et al., 2015), highlighting its essential function in the development and maintenance of these neurons. In our study, we overexpressed Activin A specifically under the human synapsin (hSyn) promoter, which drives expression primarily in neurons(Kügler et al., 2003). While neurons are believed to be the main cells secreting Activin A, it is also expressed in glial cells, and their role in the context of HD remains unclear. It is possible that impaired glial-derived Activin A may contribute to neuronal health and survival in HD, either through direct trophic support or through interactions with neuronal activity-dependent Activin A signaling.

The second potential pathway involves reduction of eNMDAR-mediated currents. In our study, we demonstrated that striatal eNMDAR-mediated currents are increased in 6- to 8-month-old YAC128 mice, consistent with previous reports at other ages and in other mouse models(Botelho et al., 2014; Kovalenko et al., 2018; Milnerwood et al., 2010b; Plotkin et al., 2014). It has been shown that glutamate spillover can trigger aberrant spike-timing-dependent plasticity (STDP) within the striatum, which may manifest as behavioral changes(Boender et al., 2021). Notably, cortico-striatal STDP-mediated long-term potentiation (t-LTP) is dependent on postsynaptic NMDARs, and more precisely, the balance between GluN2A- and GluN2B-containing NMDARs shapes STDP(Li and Pozzo-Miller, 2019). Restoring the balance between synaptic and GluN2B-enriched extrasynaptic NMDARs by Activin A expression may be crucial in re-establishing proper signal-to-noise ratios and neural communication within the cortico-striatal network, ultimately improving motor function in HD.

Interestingly, we did not find a sex difference between male and female WT and YAC128 mice in our rotarod data. There are reports, however, of sex differences in the HD population as well as in some rodent studies. Among HD patients, women have more severe motor symptoms, and men’s loss of functional abilities are more correlated with cognitive deficits(Zielonka et al., 2018). In rodents, males exhibit motor incoordination earlier and more severely than females(Cao et al., 2018; Zarringhalam et al., 2012), along with stronger effects of HD gene expression on circadian rhythm(Kuljis et al., 2016), and earlier changes in cortico-striatal circuit function and striatal neurochemistry in males(Padovan-Neto et al., 2019). In rodents, hormonal factors account for some of these sex differences(Bode et al., 2008; Hsu et al., 2011); estrogen has been shown to correlate with higher levels of DARPP32, as well as protecting neurons against apoptosis and influencing motor coordination (Nuzzo and Marino, 2016). However, there is minimal work done on the effect of sex in the YAC128 mouse line. In fact, a longitudinal study examining various neurochemical, structural, and volumetric changes in the YAC128 brain showed no sex difference at 3, 6, 9, and 12 months of age(Petrella et al., 2018). There may be sex differences on the rotarod task that appear at a different age; a dedicated study is needed to examine both male and female YAC128 mice at different time points.

### Overexpression of Activin A normalizes eNMDAR currents and restores membrane capacitance and sEPSC frequency in YAC128 MSNs

The electrophysiological findings, specifically the normalization of eNMDAR currents, membrane capacitance, and sEPSC frequency in YAC128 MSNs overexpressing Activin A, provide valuable insights into the potential benefits of Activin A overexpression in HD. In addition to its beneficial effects on circuit function and motor behavior, normalizing the balance between synaptic and extrasynaptic NMDAR activity can be neuroprotective, since eNMDAR signaling has been shown to suppress cell survival pathways by inhibiting the phosphorylation of cAMP response element-binding protein (CREB) and preventing activation of the extracellular signal-regulated kinase (ERK) pathways(Hardingham et al., 2002; Léveillé et al., 2008). In fact, we have previously reported reduced nuclear CREB activation in the striatum of YAC128 mice at both one and four months of age(Milnerwood et al., 2010b). Our findings regarding the normalization of eNMDAR currents in HD mice by Activin A overexpression align well with previous research on the FDA-approved eNMDAR antagonist memantine. Low-dose memantine, possibly due to its unique pharmacodynamics(Parsons et al., 2007), has been shown to preferentially inhibit detrimental eNMDAR activity while maintaining the beneficial synaptic signaling(Léveillé et al., 2008; Okamoto et al., 2009; Papadia et al., 2008). This selective inhibition effectively mitigated motor learning deficits in early-stage and neuropathological changes in later-stage disease in YAC128 mice(Milnerwood et al., 2010b; Okamoto et al., 2009). Our current study on Activin A adds to this body of research by highlighting the importance of targeting eNMDAR activity in developing therapeutic strategies for HD. However, to maintain long-lasting therapeutic effects, sustained treatment may be necessary. Future research should aim to address the limitations of the *in vivo* gene therapy experiments and explore alternative ways to target eNMDARs, In this regard, disruption of coupling between eNMDAR and the transient receptor potential cation channel subfamily M member 4 (TRPM4)(Yan et al., 2020) with a small molecule could be a promising approach.

In addition to the normalization of eNMDAR currents, our electrophysiological analyses revealed an increase in membrane capacitance and sEPSC frequency in YAC128 MSNs overexpressing Activin A, to levels similar to WT MSNs. The reduced capacitance together with reduced sEPSC frequency argues for reduced spine density(Buren et al., 2016) and cortico-striatal connectivity, which appears to be reversed by Activin A.

### Conclusion

Together, our findings underscore the potential for development of innovative therapeutic approaches targeting Activin A, specifically focusing on the modulation of eNMDARs, which may ultimately contribute to alleviating the debilitating symptoms and slowing the progression of HD. Future research in this area should focus on understanding the molecular mechanisms by which Activin A modulates eNMDAR currents, as well as spine density and synaptic transmission, to identify additional targets within these pathways. Longitudinal studies examining the long-term impact of Activin A treatment via systemic administration, potentially through nasal delivery as described previously(Buchthal et al., 2018), on disease progression in HD mouse models and, ultimately, in human patients are essential for evaluating the clinical potential of this approach.

## Acknowledgments

We are grateful To Dr. Lily Zhang for assistance with neuronal culture preparation, mouse genotyping and surgeries and to Dr. Jing Yan, Heidelberg University, for production of recombinant adeno-associated viruses. We acknowledge funding from the Canadian Institutes of Health Research (CIHR) PJT 178043 to LAR, Vanier Award to WN, UBC 1-year Fellowship to DR, CIHR Canada Graduate Scholarship-Masters to JC, and Hereditary Disease Foundation Fellowship to JM. Funding to HB was from the European Research Council (ERC) Advanced Grant (233034) and Deutsche Forschungsgemeinschaft (DFG) FOR 2289.

## Author contributions

Conceptualization, W.B.N., D.R., J.M., and L.A.R.; Methodology, W.B.N., D.R., J.M., and L.A.R.; Investigation, W.B.N., D.R., J.C., J.O., J.M., and M.D.S.; Resources, H.B., and L.A.R.; Writing – Original Draft, D.R., W.B.N., and L.A.R.; Writing – Review & Editing, W.B.N., D.R., J.C., J.O., J.M., M.D.S., D.L., H.B., and L.A.R.; Funding Acquisition, L.A.R.; Resources, L.A.R., and H.B.; Supervision, L.A.R., J.M., M.D.S., and H.B.

## Declaration of interests

The authors declare no competing interests.

## Materials and Methods

### Generation and Nucleofection of Cortical-Striatal Cultures

Cortical and striatal neurons were isolated from E17-18 mouse pups from FVB/N WT and homozygous YAC128 (line 55) mice colonies, bred and maintained according to the guidelines of the Canadian Council on Animal Care (protocol A21-0276 and A23-0083). This isolation procedure is based on established methods(Milnerwood et al., 2012).

One million cortical cells and one million striatal cells were suspended in 6 mL D-minimum essential medium (DMEM, GIBCO) with 10% foetal bovine serum (DMEM+). The striatal cells were then subjected to electroporation (AMAXA nucleofector I: program 05) in a buffer (Mirus Bio) along with plasmid (2ug plasmid per 2 million cells). For experiments focused on GluN2B surface/internal imaging, the neurons were transfected with the GluN2B-YFP plasmid (Syn.YFP-GluN2B; gifted from A.M. Craig, University of British Columbia; originally from M. Sheng, Genentech, San Francisco).

Post-electroporation, the striatal cells were combined with non-transfected cortical cells and plated at a density of 1.125×10^5^ cells per cm^2^ in 12-well plates, containing poly-d-lysine (PDL) coated coverslips. After three hours, the DMEM+ was replaced with 0.5 mL of neurobasal medium (NBM: 2% B27, Invitrogen; penicillin/streptomycin; 2 mM α-glutamine; in neurobasal medium A, GIBCO). An additional 0.5 mL/well of NBM was added at 3 days in vitro (DIV), and then half of the culture’s NBM was replaced at DIV10. For ELISA assays, NBM lacking a pH indicator (GIBCO) was used to prevent color interference with the Activin A ELISA results.

### Activin A ELISA and Chimeric Cultures

Activin A, a secreted protein, can be quantified in the culture media. For pure cortical-striatal cultures derived from WT and YAC128 mice, media was sampled at three distinct developmental stages - DIV 4, 10, and 21 - and subsequently stored at -80°C for further analysis. Activin A levels were measured using an ELISA kit (Human/Mouse/Rat Activin A Quantikine ELISA Kit, R&D Systems, DAC00B).

To correct for cell density variation, total protein concentration was measured using a protein assay (DC™ Protein Assay Kit II, BIO-RAD, 5000112). Activin A measurements were normalized to total protein levels, and DIV 10 and DIV 21 measurements were further normalized to the baseline taken at DIV 4.

In the case of chimeric cultures, which combine cortical and striatal neurons from both WT and YAC128 mice, media was sampled at two time points, DIV 4 and DIV 21. This approach enabled the examination of both cell-autonomous and non-cell-autonomous effects of the YAC128 genotype on Activin A release. The genotype of the source neurons could be associated with the respective changes in Activin A levels. These cultures were maintained and sampled in a similar manner as the non-chimeric cultures.

### GluN2B Surface/Internal Imaging

We used the surface to internal ratio of GluN2B as a proxy for relative extrasynaptic GluN2B-containing NMDA receptor expression(Milnerwood et al., 2012). To perform this, MSNs from cortical-striatal co-cultures were transfected with a GluN2B-YFP construct and seeded on coverslips (refer to the section on: Generation and Nucleofection of Cortical-Striatal Cultures). At DIV21, cells were stained for 10 minutes at 37°C with a chicken anti-green fluorescent protein (GFP) antibody (cross-reactive with YFP; ab13970, AbCam) diluted 1:1000 in neurobasal medium to stain for surface GluN2B. Afterward, cells were fixed for 15 minutes at room temperature in a solution of 4% paraformaldehyde (PFA) and 2% sucrose.

Cells were then washed with phosphate-buffered saline (PBS) and exposed to a secondary antibody (Alexa 488 anti-chicken, A11039, Molecular Probes, diluted 1:1000 in PBS) for 1 hour at room temperature, shielded from light and agitated gently. After the secondary antibody treatment, cells were washed with PBS containing 0.03% Triton X-100 to permeabilize the plasma membrane, and subsequently stained with chicken anti-GFP antibody (ab13970, AbCam, diluted 1:1000 in PBST) for another hour at room temperature, shielded from light and agitated gently to stain for internal GluN2B.

Finally, cells were stained with either anti-chicken Alexa 568 (red fluorescent; A21069, Molecular Probes) or anti-chicken AMCA (blue fluorescent; 706-155-148, Jackson Laboratories) conjugated secondary antibodies (diluted 1:1000) for 1 hour at room temperature and protected from the light on a shaker. The coverslips were washed with PBST and mounted onto slides using fluoromount G (Southern Biotech).

Imaging was conducted on the Zeiss Axio Observer Z1, focusing on the soma and dendritic region up to a 70 μm radius using a 63x objective to optimize the signal-to-noise ratio.

For the GluN2B surface/internal images, three regions of interest (ROIs) were manually placed on different secondary or tertiary dendrites, and a rolling ball filter was applied to subtract the background. We computed the ratio of the surface GluN2B intensity divided by the internal GluN2B intensity for each image. All images were normalized to the control wild type (WT) average within each culture batch. To ensure consistent normalization within culture batches, all cells from a single culture batch (both WT and YAC128 plated at the same time) were immune-stained simultaneously and imaged on the same day using the same excitation light intensities and exposure times per channel.

The data was then analyzed using a two-way ANOVA to explore both genotype and treatment effects. We used individual cells rather than culture batch averages as biological replicates for a more granular understanding of the effects.

### Culture rAAV Treatments

Considering Activin A’s ability to reduce the toxic eNMDAR calcium influx(Lau et al., 2015), we evaluated its impact on surface GluN2B expression through differential staining of surface and internal GluN2B. DIV 7 cortico-striatal co-cultures were treated with addition to the medium of either: i) An Activin A recombinant adeno-associated virus (rAAV) (rAAV-hSyn-inhba-HA/hSyn-tdimer, from H. Bading, Heidelberg U) at a concentration of approximately 10^9^ viral particles/mL; or ii) As a control, we used nuclear localized mCherry rAAV (rAAV-CMV/CBA-mCherry NLSr, from H. Bading, Heidelberg U). By comparing these two treatment conditions, we aimed to directly assess the impact of Activin A on the surface expression of extrasynaptic GluN2B-containing eNMDARs in our neuronal cultures.

### Stereotaxic Animal Surgeries and rAAV Injections

To investigate the therapeutic impacts of Activin A, we performed bilateral rAAV injections into the dorsal striatum of mice at 1.5 months of age. Lidocaine (50 μL) was administered subcutaneously at the surgical site for local anesthesia. During the procedure, mice were anesthetized with isoflurane (initially at 4%, then maintained at 1.5%) and received subcutaneous injections of buprenorphine (0.05 mg/kg) and meloxicam (5 mg/kg at 1 mL/kg) for pain relief. We also applied an ophthalmic ointment to both eyes post-anesthesia.

Before making the incision, we shaved and disinfected the mouse’s head with ethanol and betadine. A 0.5 cm medial (anterior-posterior) incision was then made to gain access to the skull. We drilled holes in the skull at the coordinates 2.25 mm lateral and 0.75 mm rostral relative to the bregma using a dental drill.

We used a stereotaxic injector (Fofia ZS006) to inject 1 μL of the rAAV at a rate of 2 nL/s into both hemispheres at the aforementioned coordinates and at a depth of 3.00 mm. A manipulator (Sutter Instruments) was used for accurate distance measurements. We allowed the injector to stay at the injection site for 10 minutes post-injection to facilitate passive diffusion of the rAAV.

The rAAVs injected were either an Activin A rAAV (rAAV-hSyn-inhba-HA/hSyn-tdimer, from H. Bading, Heidelberg U) at a concentration of 5 x 10^9^ viral count/mL, or a control mCherry rAAV (rAAV-CMV/CBA-mCherry NLSr, from H. Bading, Heidelberg U) at the same concentration.

After completing the injection, we sutured the incision site and allowed the mouse to recover before returning it to its cage. We then carried out daily monitoring for the next five days to ensure proper recovery.

### Rotarod Performance Test

The accelerating rotarod (Ugo Basile) was utilized to evaluate motor learning and coordination in mice at 6 months of age. This task involves a programmable rotating rod that the mice must run atop to avoid falling, with the rod’s speed gradually increasing from 5 to 40 RPM over a 300-second period.

The primary metric for this task was the latency to fall from the rod. In addition to actual falls, a “fall” was also recorded if the mouse clung to the rod and performed a full rotation. If a mouse successfully remained on the rod for the full 300 seconds, the trial was concluded and the latency to fall was recorded as 300 seconds.

Each mouse underwent three trials per day over four consecutive days, with each trial separated by a two-hour inter-trial interval. All trials were done during the day (8am-1pm) in a regular 12h light/dark cycle animal housing facility. The daily average latency to fall was computed by taking the mean of the three trials performed on that day.

### Brain Slice Preparation

Mice were anaesthetised using isoflurane and subsequently decapitated. The brain was swiftly removed, and the hemispheres were separated by bisecting along the midline. Sagittal slices of each hemisphere, with a thickness of 250 μm, were prepared with a vibratome (Leica) in ice-cold artificial cerebrospinal fluid (aCSF).

Each slice was then transferred to aCSF maintained at 37°C for a 30-minute recovery period, followed by an additional 30 minutes at room temperature prior to commencing electrophysiological experiments.

The ice-cold aCSF used for slicing (low-calcium) comprised the following components (in mM): 125 NaCl, 2.5 KCl, 25 NaHCO3, 1.25 NaH2PO4, 0.5 CaCl2, 2.5 MgCl2, and 10 glucose. The aCSF used during recovery and experiments contained (in mM): 125 NaCl, 2.5 KCl, 25 NaHCO3, 1.25 NaH2PO4, 2 CaCl2, 1 MgCl2, and 10 glucose. The pH of the aCSF was maintained between 7.3 and 7.4, and its osmolarity was set at 310 Osm/L.

Throughout the slicing and recovery processes, as well as during experiments, the aCSF was continuously oxygenated with carbogen (95% O_2_ and 5% CO_2_). Once transferred to the recording chamber, slices were continuously perfused with room temperature aCSF, which also contained picrotoxin (50 μM; Tocris Bioscience) to inhibit GABA-A receptors.

### Electrophysiology

Whole-cell patch clamp recordings were carried out in voltage-clamp mode. The recordings were obtained using an amplifier (MultiClamp 700A Molecular Devices) and a digitizer (Molecular Devices 1550B). The data were digitized at 20 kHz and captured with a low pass 1 kHz bessel filter.

Electrode pipettes were pulled from borosilicate glass capillaries with the aid of a micropipette puller (Narishige International) for both the recording and stimulating electrodes. The intracellular solution used was cesium-based, as cesium effectively blocks potassium channels, thereby facilitating a superior voltage-clamp. The intracellular solution contained the following (in mM): 130 cesium methanesulfonate, 5 CsCl, 4 NaCl, 1 MgCl2, 5 EGTA, 10 HEPES, 5 QX-314 chloride, 5 MgATP, 0.5 MgGTP, and 10 sodium phosphocreatine.

The pH of the intracellular solution was maintained between 7.25 and 7.3, and its osmolarity was 290 (±3) mOsm/L. Only medium spiny neurons (MSNs) with an initial access resistance of less than 17 MΩ, and less than 10% change in series resistance throughout the experiment were considered for the analysis. In all cells, recording only commenced after a five-minute period following access acquisition to allow for the equilibration of the electrode solution with the intracellular space. To record spontaneous excitatory postsynaptic currents (sEPSCs), cells were voltage-clamped at -70 mV.

### eNMDAR-mediated Current Measurement

Investigation of eNMDARs was carried out by achieving a whole-cell voltage clamp, followed by the generation of stimulation trains (20 Hz for 500 ms) using a stimulating electrode positioned approximately 200 μm dorsal to the cell being recorded. The cell was voltage-clamped at +40 mV to remove the Mg^2+^ block from NMDARs, and a 50 ms voltage step to +30 mV was applied every 30 s to monitor the resistance.

To isolate NMDAR current, CNQX (10 μM, Tocris; an AMPA receptor antagonist), glycine (10 μM, Sigma; an NMDA receptor co-agonist), and LY341495 (1uM, Tocris; to inhibit presynaptic metabotropic glutamate receptors) were utilised. DL-TBOA (10 μM, Tocris), a glutamate transport inhibitor, was employed to cause glutamate spillover beyond the synapse into extrasynaptic sites, activating eNMDARs. The inclusion of LY341495 in the TBOA solution was critical to prevent glutamate spillover-induced activation of presynaptic mGluR2/3 receptors, reducing glutamate release.

The analysis of electrophysiology data was conducted using Clampfit 10.7 (Molecular Devices). The area under the 10th peak was measured and then normalized to the 10th peak amplitude. The decay of the 10th peak is representative of glutamate spillover and eNMDAR activation. Therefore, by normalizing to the 10th peak amplitude, we can assess the relative contribution of eNMDAR activation. This normalization process was essential to control for variability across different cells and experiments in terms of stimulation amplitude, enabling comparable and reliable assessment of the impact of glutamate spillover on activation of eNMDARs.

## Notes

### Competing Interest Statement

The authors have declared no competing interest.

