## Supplementary material for "Activin A targets extrasynaptic NMDA receptors to improve neuronal and behavioral deficits in a mouse model of Huntington disease": Figure 3 Supplemental Figure

**Figure 3 - Supplement**

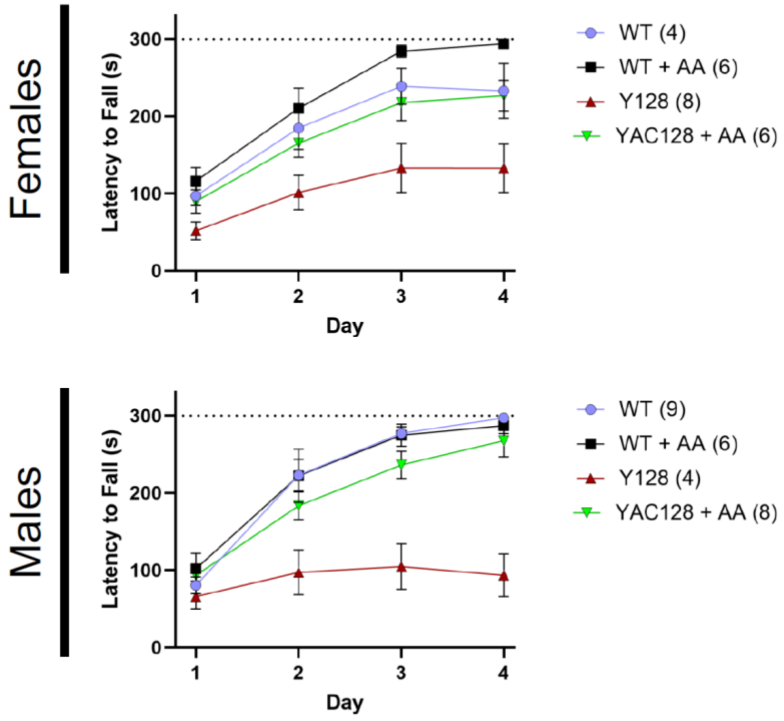


**Figure 3.s. Activin A overexpression increases the ability of YAC128 mice to learn the rotarod task for females and males**. Latency to fall day averages of WT and YAC128 female (*top*) and male (*bottom*) mice at 6 months of age treated either with Activin A rAAV or a control mCherry rAAV. Data were analysed with a two-way ANOVA (***Female*** p-values: genotype = 0.0008, Days = <0.0001, interaction = 0.1077; ***Male*** p-values: genotype = <0.0001, Days = <0.0001, interaction = <0.0001). Data are represented as mean ± SEM (AA = Activin A, WT = Wild-type, Y128 = YAC128). Number in parentheses indicates number of animals.
